# Mechanisms of self-assembly and fibrillization of the prion-like domains

**DOI:** 10.1101/065631

**Authors:** Yimei Lu, Liangzhong Lim, Yanming Tan, Lu Wang, Jianxing Song

**Author notes:** The first three authors contribute equally.

## Abstract

The mechanism of the self-assembly and fibrillization of the prion-like domains lies at the heart of their physiology and pathology. Here with the same methods previously established, we aimed to further decipher the mechanism by characterizing two prion-like sequences with the electrostatic properties very different from that of the full-length TDP-43 prion-like domain with a very basic pI value: namely the C-half of the TDP-43 prion-like domain only abundant in Gly, Ser, Asn and Gln with a pI of ~6.3, and the FUS prion-like domain enriched with Gly, Ser, Gln and Tyr with a pI of ~3.5. Interestingly, the C-half with the TDP-43 unique hydrophobic region removed is no longer able to form insoluble aggregates/fibrils but still capable of self-assembling into the reversible hydrogel with cross-β structures, despite being much slower than the full-length. On the other hand, the FUS prion-like domain rapidly self-assembles into the reversible hydrogel with cross-β fibrillar structures in 1 mM phosphate buffer at pH 6.8 but its self-assembly becomes very slow in 50 mM MES buffer at pH 5.5. Our study reveals that despite having completely different electrostatic properties, the full-length and C-half of the TDP-43 prion-like domain, as well as FUS prion-like domain all have the similar pH-dependence in self-assembly as we previously reported (Lim et al., [2016] PLOS Biol 14:e1002338). This unambiguously indicates that the self-assembly of the prion-like domains is not generally governed by the electrostatic interaction. Rather, their self-assembly and fibrillization are specified by the sequences despite being highly polar and degenerative. Furthermore, our study provides the first evidence that the formation of reversible hydrogel with cross-β structures is separable from fibrillization of the prion-like domain. Finally, our results also successfully reconcile the previous discrepancy about the conformation and mechanism of the self-assembly of the FUS prion-like domain.

## Introduction

Prions are characteristic of their amazing capacity of adopting many distinct conformations, some of which are infectious [1–5]. While the mammalian prion protein (PrP) causes devastating and infectious neurodegenerative diseases, yeast prions such as Sup35 have been revealed to confer selective advantages [2–4]. Yeast prions are characterized by low complexity (LC) sequences enriched in polar and uncharged amino acids such as Gln, Asn, Ser, Gly and Tyr. Recently 240 genes out of ~20,000 human protein-coding genes (~1.2%) were shown to contain at least a domain compositionally similar to yeast prions, thus termed prion-like domains [4]. Remarkably, human proteins containing the prion-like domains are highly overrepresented by those critically interacting with RNA and DNA through self-assembling into physiological and reversible droplet/hydrogel, which may be exaggerated into pathological and irreversible aggregates/fibrils characteristic of a variety of neurodegenerative diseases [3–10]. Therefore decoding molecular mechanisms underlying the self-assembly, aggregation and fibrillization of the prion-like domains lies at the heart of their physiology and pathology; and may offer rationales to further develop therapeutic strategies for the related diseases.

Previously, as facilitated by our previous discovery that protein aggregation can be significantly minimized by reducing salt concentrations [10–17], we showed that although the TDP-43 prion-like domain is intrinsically disordered, it has the capacity to spontaneously self-assemble into dynamic and cross-β oligomers while ALS-causing point mutations are sufficient to exaggerate the formation of the amyloid-fibers [10]. Very interestingly, we observed that the self-assembly of the TDP-43 prion-like domain is highly pH-dependent: it showed no significant oligomerization at acidic pH but started to self-assemble at neutral pH. To rationalize this, we proposed [10] that at acidic pH, the majority of the side chain atoms of Asn, Gln and Ser are involved in forming hydrogen bonds with the backbone atoms as extensively observed in both well-folded proteins as well as disordered short peptides [18–20]. However, the self-assembly will be initiated by the liberation of the Asn, Gln and Ser side chains at neutral pH to form inter-molecular “hydrogen bonds/steric zippers” as well established [3, 6, 21, 22]. As such, despite being highly sequence-degenerative and intrinsically disordered, the prion-like domains, such as human TDP-43, appear to represent a subgroup of the intrinsically disordered proteins that can achieve extremely high specificity in the self-assembly on the basis of the global networks constituted by sequence-dependent intramolecular hydrogen-bonding/interactions [10]. Very recently, however, it was proposed in a formal comment that the pH-dependent assembly of the TDP-43 prion-like domain we observed [10] can be simply interpreted as “Electrostatic Repulsion Governs TDP-43 C-terminal Domain Aggregation” [23].

To adequately clarify this issue, here we present the new experimental results on the C-half of the TDP-43 prion-like domain over residues 342-414 (Fig 1A) and the FUS prion-like domain over residues 1-165 (Fig 1B). Based on our previous NMR characterization [10], the TDP-43 prion-like domain radically differs from other prion-like domains in having a unique hydrophobic region over Met307-Gln344, which transforms into a well-folded Ωloop-helix motif upon being embedded in membrane environments. So here we focused only on the 73-residue C-half of the TDP-43 prion-like domain with a pI of ~6.3, which is mostly enriched in Asn, Gln, Ser and Gly (Fig 1C). Furthermore, we also studied the FUS prion-like domain with a pI of ~3.5, which has much higher contents of Gln and Tyr than those of the C-half of the TDP-43 prion-like domain (Fig 1D).

**Fig 1.**
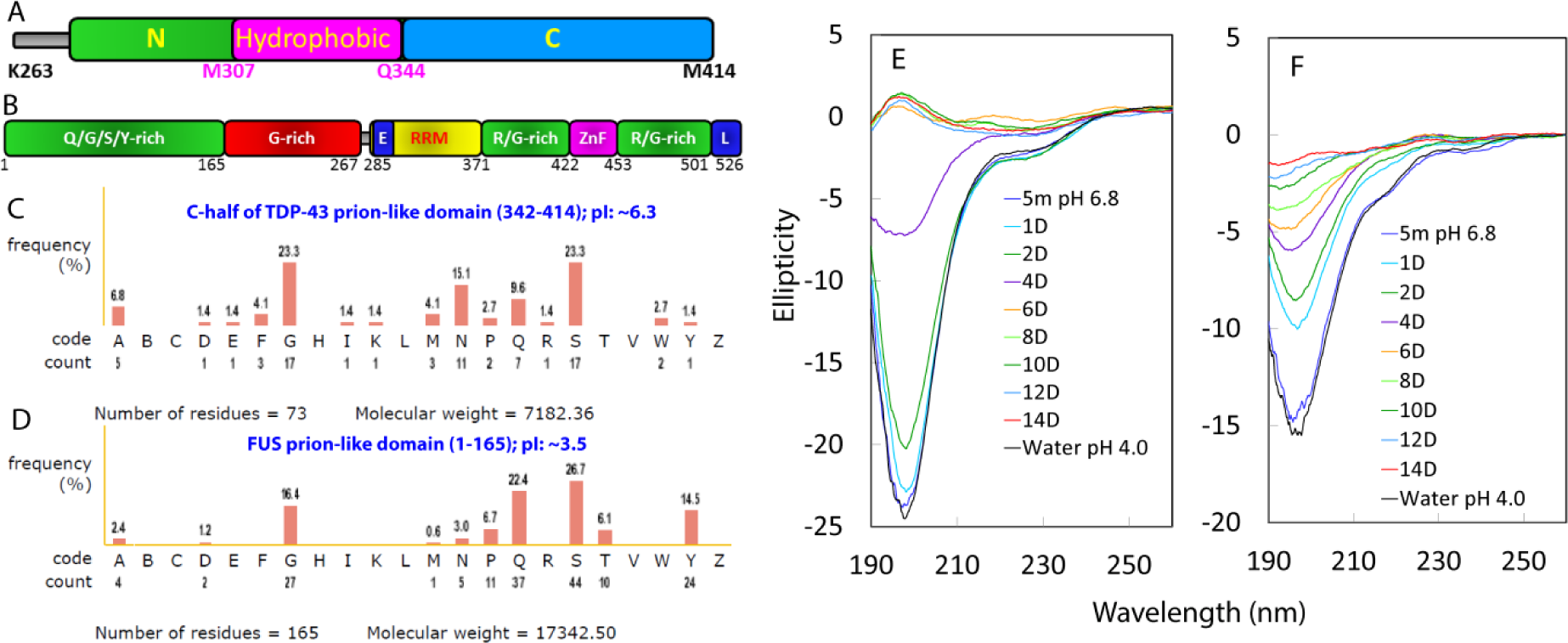
Domain organizations and bioinformatics analysis. Domain organizations of the prion-like domain of TDP-43 (A); and FUS protein (B). Amino acid compositions of the C-half of the TDP-43 prion-like domain (C); and FUS prion-like domain (D). Far-UV CD spectra of the C-half (E); and FUS prion-like domain (F) at a protein concentration of 40 µM in Milli-Q water (pH 4.0) as well as at different time points of the incubation in 1 mM phosphate buffer (pH 6.8).

## Results

### The self-assembly of the C-half

In Milli-Q water (pH 4.0), the C-half has a predominantly disordered conformation very similar to that of the full-length [10], as evident from its CD (Fig 1E). Furthermore, the C-half residues adopt the solution conformation very similar to that of the corresponding region of the full-length TDP-43 prion-like domain, as judged from the fact that most of its HSQC peaks are superimposable to those of the corresponding residues of the full-length (Fig 2A). Like the full-length, it showed no detectable self-assembly or aggregation for several months at high protein concentrations (~300 µM) in Milli-Q water (pH 4.0). However, upon diluting into 1 mM phosphate buffer (pH 6.8), ~60% HSQC peaks disappeared, particularly over the N-terminal Gln-rich region (Fig 2B-2D). On the other hand, its 1D NMR peaks for the methyl protons at pH 6.8 remained very similar to those in Milli-Q water (Fig 2E), suggesting that the disappearance of the HSQC peaks is mostly due to the rapid exchange of these exposed backbone amide protons with the bulk water at pH 6.8.

**Fig 2.**
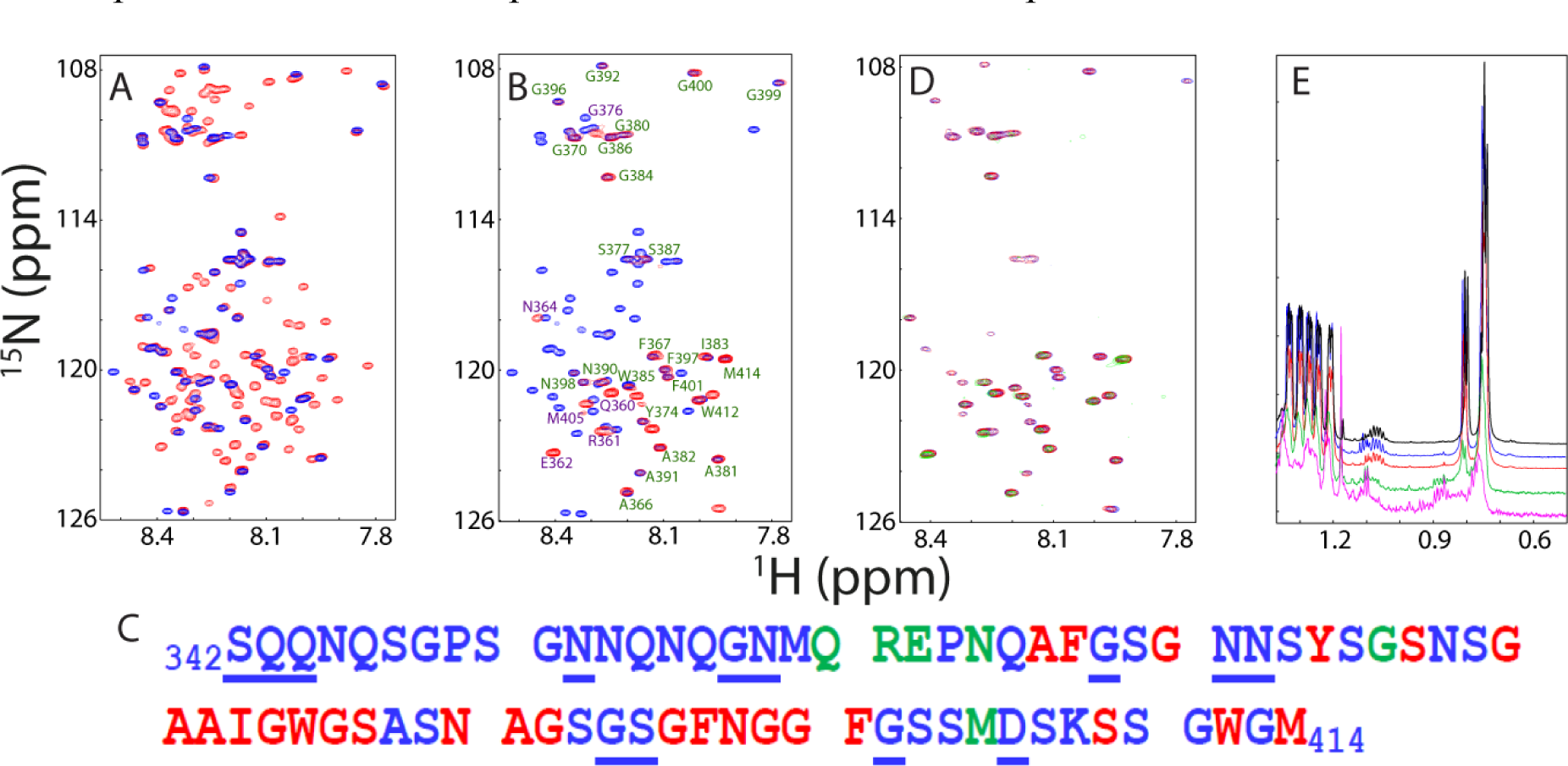
NMR characterization of the C-half. (A) Superimposition of HSQC spectra of the full-length TDP-43 prion-like domain (red) and C-half (blue) at a protein concentration of 40 µM in Milli-Q water at pH 4.0. (B) Superimposition of HSQC spectra of the C-half in Milli-Q water at pH 4.0 (blue) and that immediately diluted into 1 mM phosphate buffer at pH 6.8 (red) at a protein concentration of 40 µM. (C) The C-half sequence with the residues unassigned underlined; those retained and superimposable to those in Milli-Q water at pH 4.0 (red), retained but significantly shifted (green) upon immediately dilution into 1 mM phosphate buffer at pH 6.8. (D) Superimposition of HSQC spectra of the C-half at a protein concentration of 40 µM immediately diluted into 1 mM phosphate buffer at pH 6.8 (blue); at day 6 (red); day 10 (green). (E) One-dimensional NMR spectra over 0.5-1.35 ppm at a protein concentration of 40 µM acquired in Milli-Q water at pH 4.0 (black); immediately diluted into 1 mM phosphate buffer at pH 6.8 (blue), 6 day (red), 10 day (green) and 14 day (purple).

Like the full-length, at pH 6.8 the C-half was still capable of gradually self-assembling into oligomers rich in β-structure as monitored by CD, fluorescence and EM. As seen in Fig 1E, in the first 4 days, the absolute values of the CD signal at ~198 nm continuously reduced, indicating a spontaneous and gradual self-assembly. A transition occurred from 4 to 6 day. After 6 day, the CD spectra remained largely unchanged, which are typical of the soluble cross-β oligomers such as formed by the wild type TDP-43 prion-like domain [10]. Consistent with the CD results, both HSQC (Fig 2D) and 1D (Fig 2E) spectra at day 6 remained highly similar, only with a slight reduction of peak intensity, as compared to those collected at 15 min after dilution into 1 mM phosphate buffer at pH 6.8. However, in 10 day, the intensity of the HSQC and 1D peaks became significantly reduced and at day 14, almost all HSQC peaks became too broad to be detected. Furthermore, after 10 days, the sample became dynamic hydrogel at 4 °C but could be converted back to liquid at room temperature or/and by shaking as previously observed on the FUS [6] and full-length TDP-43 prion-like domains [10].

Interestingly, the time-lapse change of the intrinsic UV fluorescence revealed a multi-phase process of the self-assembly of the C-half which contains two Trp residues: Trp385 and Trp412 (Fig 2C). As seen in the upper panel of Fig 3A, similar to what was observed on the full-length in Milli-Q water at pH 4.0 [10], the C-half has an intrinsic UV fluorescence spectrum with the emission maximum of 352 nm, which implies that both Trp residues are highly exposed as in the full-length. Immediately after dilution into the 1 mM phosphate buffer at pH 6.8, a slight increase of the intensity was observed. However, within the first 2 days of the incubation, the intensity reduced and the emission maximum became largely blue-shifted to ~332 nm. Amazingly, from day 4 to day 6, the spectra suddenly have two emission maxima: one at ~335 nm and another at ~385 nm. This implies that in this period of the self-assembly, two Trp residues may experience different chemical environments. After day 8, the intensity further reduced and only one emission maximum was observed at ~324 nm, suggesting that both Trp residues became highly buried.

**Fig 3.**
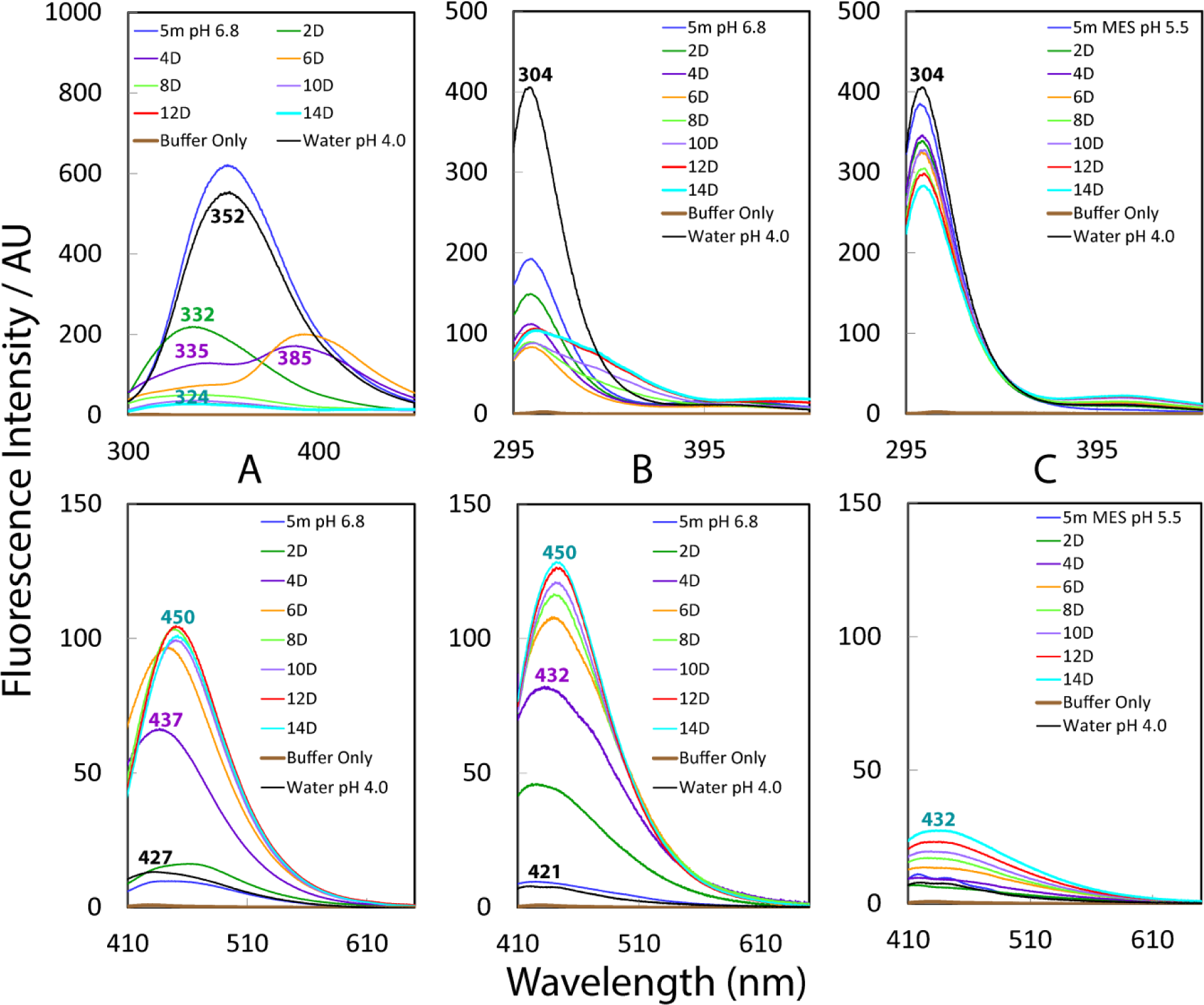
Fluorescence characterization. Emission spectra of the intrinsic UV (upper panel) and visible (lower panel) fluorescence at different time points of the incubation at a protein concentration of 40 µM respectively for the C-half of the TDP-43 prion-like domain in 1 mM phosphate buffer at pH 6.8 (A), FUS prion-like domain in 1 mM phosphate buffer at pH 6.8(B) and FUS prion-like domain in 50 mM MES buffer at pH 5.5 (C).

Recently, an intrinsic visible fluorescence has been established to be a signature for the cross-β structures, which is independent of the presence of aromatic residues but have its origin in the formation of aligned hydrogen bond networks involved in the backbone C = O and N-H atom groups of peptide bonds [10, 24, 25]. Like the full-length, the C-half samples both in Milli-Q water at pH 4.0 and immediately diluted in 1 mM phosphate buffer at pH 6.8 already have the intrinsic visible fluorescence but its intensity (12.8) is lower than that (25.1) of the full-length, while the emission maximum (~432 nm) is also less red-shifted than that of the full-length (~446 nm) (lower panel of Fig 3A). Nevertheless, with the time lapse, the intensity continuously increased and consequently after 12 days, it reached the highest (104.3) and the emission maximum red-shifted to ~450 nm. By contrast, under the same conditions, only after 2 days, the intensity of the full-length reached the highest (~62.5) with the emission maximum red-shifted to ~450 nm. This indicate that: 1) the self-assembly of the C-half is much slower than that of the full-length; 2) the C-half could still form reversible hydrogel even with the higher content of cross-β structures.

Unexpectedly, however, very different from the full-length, as imaged by EM, the C-half failed to form any insoluble aggregates/fibrillar structures even after 2 weeks of the incubation (Fig 4A). By contrast, under the exact same conditions, both 1D and HSQC peaks of the full-length became very broad after 24 hr, and further incubation led to formation of insoluble fibrils [10]. The fact that the C-half with a pI of ~6.3 self-assembled into the cross-β hydrogel much more slowly than the full-length with a pI of 10.9, clearly indicates that the self-assembly of the TDP-43 prion-like domain is not simply governed by electrostatic interaction.

**Fig 4.**
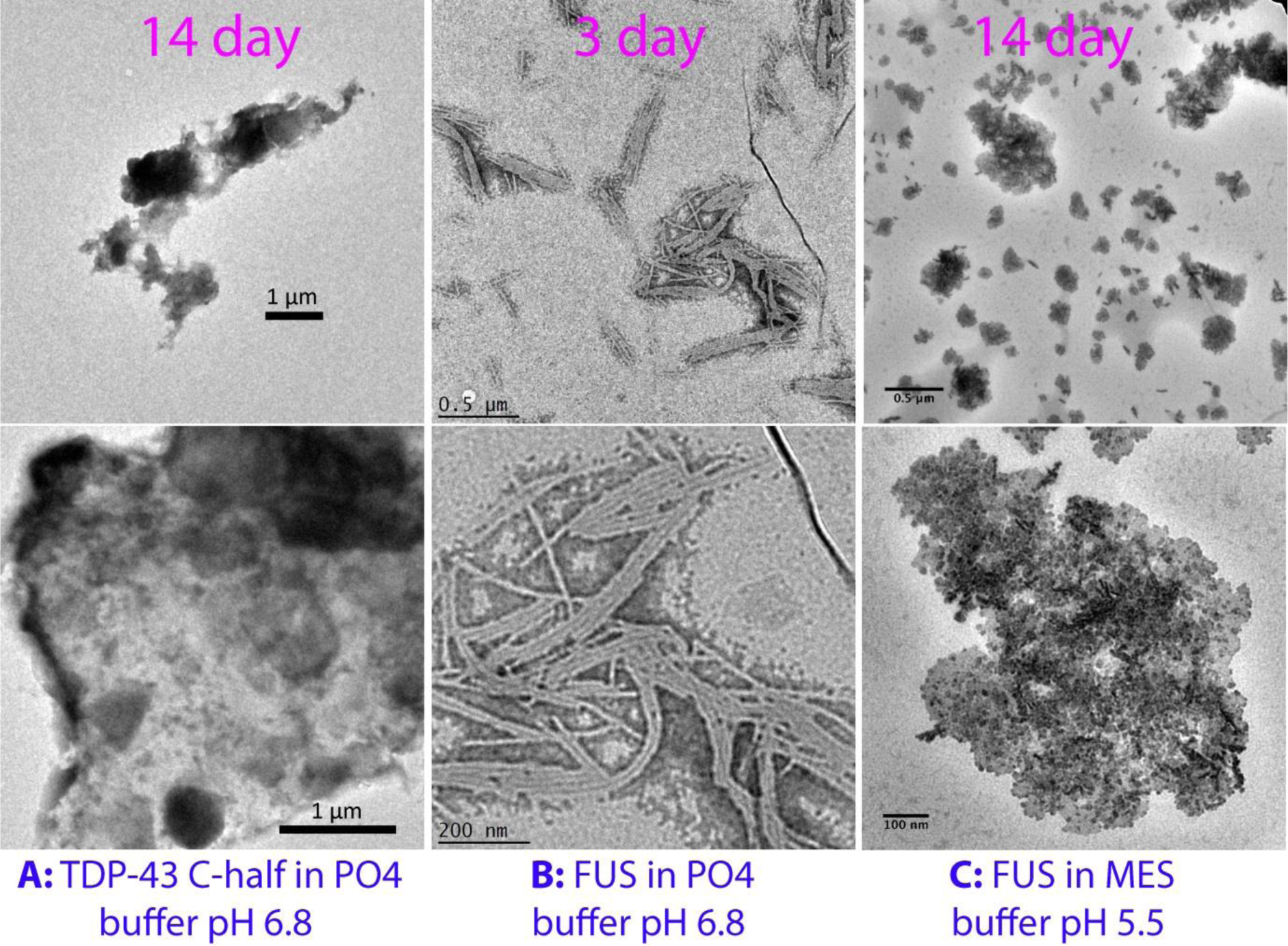
EM imaging. EM images of the samples of the C-half incubated for 14 days in 1 mM phosphate buffer at pH 6.8 (A); of FUS prion-like domain incubated for 3 days in 1 mM phosphate buffer at pH6.8 (B); and of FUS prion-like domain incubated for 14 days in 50 mM MES buffer at pH 5.5
(C).

### The self-assembly and fibrillization of the FUS prion-like domain

Intriguingly, despite having an acidic pI value, the FUS prion-like domain also showed no detectable self-assembly at high concentrations (~400 µM) in Milli-Q water (pH 4.0). It has a highly disordered conformation in Milli-Q water (pH 4.0), as indicated by its CD spectrum with the maximum negative signal at ~196 nm but no positive signal (Fig 1F). Furthermore, we have acquired one-dimensional and two-dimensional NMR HSQC spectra for the FUS prion-like domain at a protein concentration of 40 µM in Milli-Q water at pH 4.0, 1 mM phosphate buffer at pH 5.0 and 6.8 respectively, as well as in 50 mM MES (N-morpholinoethanesulfonic acid) buffer at pH 5.5 respectively (Fig 5A-5D). In Milli-Q water at pH 4.0, the FUS prion-like domain has a narrowly-dispersed HSQC spectrum with only the^1^H dispersion of 0.95 ppm and ^15^N dispersion of 17.3 ppm (Fig 5A); and has no very up-field peaks with negative chemical shifts in its 1D spectrum (Fig 5B). This indicates that the FUS prion-like domain has no tight tertiary packing. However, very different from what we observed on the TDP-43 prion-like domain [10], even in 1 mM phosphate buffer at pH 5.0, a portion of HSQC peaks completely disappeared and many HSQC peaks shifted (Fig 5A), clearly indicating that these amide protons might be highly exposed to the bulk solvent. On the other hand, its 1D spectrum over 1-2 ppm at pH 5.0 is still very similar to that in Milli-Q water at pH 4.0 (Fig 5B), indicating that no significantly self-assembly occurred shortly at pH 5.0. Interestingly, the majority of its HSQC peaks at pH 4.0 and pH 5.0 (Fig 5A) are superimposable to those in 50 mM MES buffer at pH 5.5 (Fig 5C), under which the same FUS prion-like domain has been characterized to exist as a highly disordered monomer by NMR [9].

**Fig 5.**
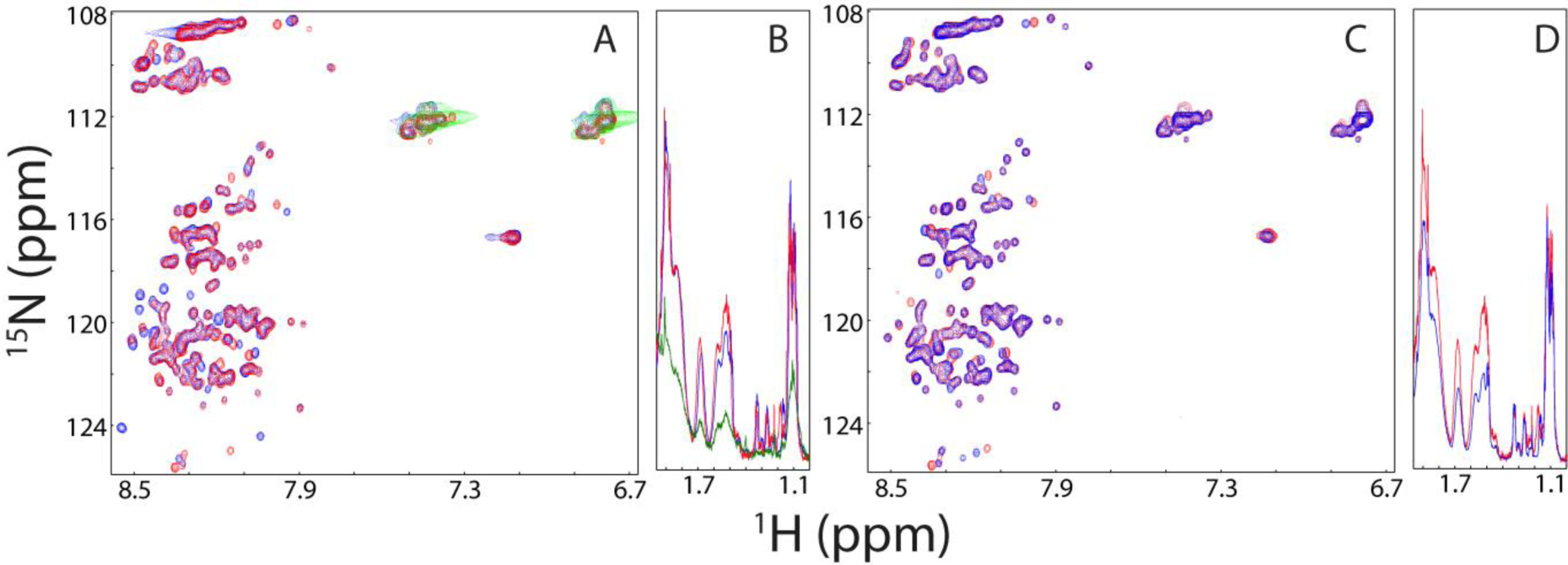
NMR characterization of the FUS prion-like domain. Superimposition of HSQC (A) and 1D ^1^H (B) NMR spectra of the FUS prion-like domain in Milli-Q water at pH 4.0 (blue); in 1 mM phosphate buffer at pH 5.0 (red); and at pH 6.8 (green). Superimposition of HSQC (C) and 1D ^1^H (D) NMR spectra of the FUS prion-like domain in 1 mM phosphate buffer at pH 5.0 (blue) and in 50 mM MES buffer at pH 5.5 (red).

However, upon diluting the FUS prion-like domain into 1 mM phosphate buffer at pH 6.8, its 1D peaks became very broad (Fig 5B), while almost all backbone HSQC peaks disappeared (Fig 5A), mostly due to rapid exchanges of amide protons with water, or/and the broadening of peaks resulting from the rapid self-assembly into large oligomers as previously characterized [10]. This is very different from the previous results with the full-length TDP-43 prion-like domains: even in 1 mM phosphate buffer at pH 6.8, the majority of HSQC peaks was still detectable and only became too broad to be detectable after 4 day at the same concentration of 40 µM. The present results clearly indicate that the FUS prion-like domain has its backbone amide protons much less protected, or/and a tendency in self-assembly much higher than TDP-43 [10].

Consistent with the above NMR observation, very large changes of the CD intensity were observed for FUS prion-like domain after 1 day (Fig 1F), suggesting its rapid self-assembly. As the FUS prion-like domain has no Trp but has 24 Tyr residues, its intrinsic UV fluorescence from Tyr has the maximum emission wavelength at ~304 nm (Upper panel of Fig 3B). Interestingly, its fluorescence intensity significantly reduced immediately upon being diluted into 1 mM phosphate buffer at pH 6.8. For the intrinsic visible fluorescence (Lower panel of Fig 3B), although FUS prion-like domain has very weak emission signal 5 min after diluting into 1 mM phosphate buffer at pH 6.8, it rapidly developed after 2 days with the emission maximum at ~425 nm and intensity of 46.2. After 4 days it further developed with the emission maximum red-shifted to ~432 nm and intensity increased to of 81.8. After 14 days it fully developed with the emission maximum red-shifted to ~450 nm and intensity increased to of 126.5, which is even much higher than that developed for the full-length TDP-43 prion-like domains [10]. The results reveal that at pH 6.8, the highly disordered FUS prion-like domain could indeed self-assemble into the hydrogel with cross-β structures. Furthermore, it forms fibrillar structures with diameter of 10-20 nm even in 3 days (Fig 4B), and no significant change of the morphology was observed even after 14 days. Our current results are completely consistent with the self-assembly of the FUS prion-like domain previously revealed by EM and other biophysical probes including X-ray diffraction [6].

We also prepared a sample of the FUS prion-like domain in 50 mM MES buffer at pH 5.5, which was previously used to NMR characterize the same FUS prion-like domain [9]. Due to the large noise from the MES buffer, the self-assembly could not be monitored by far-UV CD spectroscopy. Nevertheless, it has HSQC and 1D NMR spectra very similar to those in 1 mM phosphate buffer at pH 5.0 (Fig 5C and 5D). Furthermore, very small change in the intrinsic UV fluorescence was observed upon immediate dilution into 50 mM MES buffer at pH 5.5 (Upper panel of Fig 3C). Most remarkably, unlike in 1 mM phosphate buffer at pH
6.8 (Upper panel of Fig 3B), in 50 mM MES buffer at pH 5.5, only small changes in the intrinsic UV fluorescence have been detected with the incubation up to 14 days (Upper panel of Fig 3C). Consistent with the UV intrinsic fluorescence, only very small increases in the intrinsic visible fluorescence with minor red-shifts have been observed during incubation up to 14 days (Lower panel of Fig 3C). Moreover, the sample remained transparent and failed to form hydrogel even up to 14 days. Indeed, no fibrillar structure was detected by EM for the sample incubated up to 14 days (Figure 4C).

## Discussion

To further decipher the molecular determinant for the self-assembly and fibrillization of the prion-like domains, with the same methods we previously utilized [10], we characterized two prion-like sequences with electrostatic properties different from that of the full-length TDP-43 prion-like domain, which are only enriched in polar and uncharged residues Gly, Ser, Asn, Gln and Tyr characteristic of the prion-like domains [4]. The new results, together with previous one [10] reveal that despite the complete difference of their electrostatic properties, all three prion-like domains have the same pH-dependent self-assembly: the self-assembly is significantly inhibited at acidic pH while it is initiated at neutral pH as we reported before [10]. Strikingly, the C-half with its pI much closer to neutral pH than the full-length self-assembled much slowly than the full-length TDP-43 prion-like domain. The results together clearly indicate that the self-assembly of the prion-like domains is not generally governed by the electrostatic interactions as commented [23]. Therefore, to the best of our knowledge, the pH-dependent self-assembly of the TDP-43 and FUS prion-like domains can only be rationalized by the mechanism we previously proposed [10]. This mechanism is further supported by the observation here that the backbone amide protons of the FUS prion-like domain are much more significantly perturbed by the pH changes, as the FUS prion-like domain contains much higher content of Gln than the TDP-43 prion-like domain (Fig 1D). It has been shown that the hydrogen bonds formed between side chain and backbone atoms of Gln are less stable than those of Asn [18–20].

Our current results show for the first time that the self-assembly into dynamic hydrogels with significant cross-β structures and fibrillization are separable for the prion-like domains. This suggests that the prion-like sequences enriched in the polar and uncharged residues are able to spontaneously self-assemble into dynamic hydrogels with significant cross-β structures, but whether they can further fibrillize is highly dependent on the strength of the interactions. For example, the full-length TDP-43 prion-like domain appears to employ the hydrophobic interaction offered by the middle region, while the FUS prion-like domain utilizes the aromatic interaction provided a large number of Tyr residues to drive further fibrillization. It is highly possible that many environmental factors may act to enhance the interaction strength, thus driving the jump from dynamic hydrogels into fibrillar structures, both of which share the same cross-β structures.

Finally, our results also successfully reconcile the previous discrepancy about the mechanism by which the FUS prion-like domain drives the self-assembly: in one study conducted in neutral pH, the FUS prion-like domain has been characterized to self-assemble into dynamic hydrogel with fibrillar structures [6], while in a later NMR characterization performed in 50 mM MES buffer at pH 5.5, the FUS prion-like domain was shown to remain highly disordered even upon forming droplets active for binding [9]. Based on our present results, it seems that the relatively disordered state previously observed [9] represents a snapshot of the self-assembly at the early stages, while the dynamic hydrogel with fibrillar structures [6] reflects the structures formed at the late stages. Taken together, our results suggest that the formation of cross-β structures represents a common mechanism for the self-assembly of the prion-like domains, but whether further fibrillization occurs is mediated by the intrinsic sequence-specific features as well as environmental factors. In cells, different snapshots of the self-assembly might be all utilized for implementing the biological functions by the proteins containing the prion-like domains [6–9,26].

## Methods

### Preparation of recombinant proteins

The DNA fragments encoding the C-half of the TDP-43 prion-like domain (342-414) and FUS prion-like domain (1-165) were amplified by PCR reactions from TDP-43 and FUS cDNA and subsequently cloned into a modified vector pET28a with 6 His residues at C-terminus as we previously used for the TDP-43 prion-like domain [10]. The expression vectors were subsequently transformed into and overexpressed in *Escherichia coli* BL21 (DE3) cells (Novagen). The recombinant protein of the C-half of the TDP-43 prion-like domain was found in supernatant while FUS prion-like domain in inclusion body. As a result the TDP-43 prion-like domain was purified by a Ni^2+^-affinity column (Novagen) under native conditions; while FUS prion-like domain was purified by a Ni^2+^-affinity column (Novagen) under denaturing conditions in the presence of 8 M urea. The fractions containing the recombinant proteins were acidified by adding 10% acetic acid and subsequently purified by reverse-phase (RP) HPLC on a C4 column eluted by water-acetonitrile solvent system. The HPLC elutions containing pure recombinant proteins were lyophilized [27].

The generation of the isotope-labelled proteins for NMR studies followed a similar procedure except that the bacteria were grown in M9 medium with the addition of (^15^NH_4_)_2_SO_4_ for ^15^N labeling and (^15^NH_4_)_2_SO_4_/[^13^C]-glucose for double labelling [27]. The purity of the recombinant proteins was checked by SDS–PAGE gels and their molecular weights were verified by a Voyager STR matrix-assisted laser desorption ionization time-of-flight-mass spectrometer (Applied Biosystems). The concentration of protein samples was determined by the UV spectroscopic method in the presence of 8 M urea. Briefly, under the denaturing condition, the extinct coefficient at 280 nm of a protein can be calculated by adding up the contribution of Trp, Tyr and Cys residues.

### CD and NMR experiments

All circular dichroism (CD) experiments were performed on a Jasco J-810 spectropolarimeter equipped with a thermal controller using 1-mm path length cuvettes. Data from five independent scans were added and averaged. CD samples were prepared by diluting the concentrated samples (~300 µM) dissolved in Milli-Q water (pH 4.0) into different buffers respectively. All NMR experiments were acquired on an 800 MHz Bruker Avance spectrometer equipped with pulse field gradient units as described previously [27,28].

### Fluorescence spectral measurements

All fluorescence spectra were measured at 25 °C with a RF-5301 PC spectrophotometer (Shimadzu, Japan) as previously established [10] at different time points of the incubations at a protein concentration of 40 µM in different buffers. The rectangular fluorescence quartz cuvette has the pathlength dimension of 10 × 10 mm and the general settings are: PMT at low sensitivity and scan speed of medium speed (200 nm/min). For the intrinsic UV fluorescence, the emission spectra were measured with the excitation wavelength at 280 nm and slit widths: excitation at 5 nm and emission at 10 nm. For the intrinsic visible fluorescence, the emission spectra were measured with the excitation wavelength at 375 nm and slit widths: excitation at 20 nm and emission at 10 nm.

### Electron microscopy imaging

Incubation samples at a protein concentration of 40 µM were imaged at different time points by a TEM microscope (Jeol Jem 2010f Hrtem, Japan) operating at an accelerating voltage of 200 kV. For EM imaging, a 5 µl aliquot of the incubation or aggregate solutions was placed onto the Cu grids (coated with carbon film; 150 mesh; 3mm in diameter) and negatively stained with 5 μl of 2% neutral, phosphotungstic acid (PTA). This aliquot was allowed to settle on Cu grid for 30 sec before the excess fluid was drained away. The Cu grid was later air-dried for another 15 mins before being imaged.

## Author Contributions

Conceived and designed the experiments: JXS; Performed the experiments: YML LZL YMT LW. Analyzed the data: JXS YML LZL YMT. Wrote the paper: JXS.

## Acknowledgement

This study is supported by Ministry of Education of Singapore (MOE) Tier 2 Grants 2011-T2-1-096 and MOE2015-T2-1-111 to Jianxing Song. The funders had no role in study design, data collection and analysis, decision to publish, or preparation of the manuscript.

